# Lipid droplet surface promotes three-dimensional morphological evolution of non-rhomboidal cholesterol crystals

**DOI:** 10.1101/2024.07.18.604130

**Authors:** Hyun-Ro Lee, Seunghan Kang, Siyoung Q. Choi

**Affiliations:** Department of Chemical and Biomolecular Engineering, Korea Advanced Institute of Science and Technology (KAIST), Daejeon 34141, Republic of Korea; Advanced Battery Center, KAIST Institute for the NanoCentury, Korea Advanced Institute of Science and Technology (KAIST), Daejeon 34141, Republic of Korea

**Author notes:** These authors contributed equally to this work (Hyun-Ro Lee, Seunghan Kang).

## Abstract

Cholesterol crystals, which cause inflammation and various diseases, predominantly grow in a platy, rhomboid structure on the plasma membranes but exhibit an uneven three-dimensional architecture intracellularly. Here, we demonstrate how cholesterol crystallizes in a non-rhomboidal shape on the surface of lipid droplets and develops into three-dimensional sheet-like agglomerates using an *in vitro* lipid droplet reconstitution system with stereoscopic fluorescence imaging. Our findings reveal that interfacial cholesterol transport on the lipid droplet surface and unique lipid droplet components significantly influence the nucleation-and-growth dynamics of cholesterol crystals, leading to crystal growth in various polygonal shapes. Furthermore, cholesterol crystals readily agglomerate to form large, curved sheet structures on the confined, spherical surfaces of lipid droplets. This discovery enhances our understanding of the volumetric morphological growth of intracellular cholesterol crystals.

## Introduction

Cholesterol is essential for modulating cell signaling, maintaining cellular membrane integrity, and producing steroid hormones, vitamin D, and bile acids (*1*). However, excessive cholesterol generates water-insoluble, solid crystals that trigger mechanical tissue damage and inflammatory responses, potentially causing fatal clinical disorders such as atherosclerotic cardiovascular diseases (*2*, *3*), demyelinating disorders (*4*, *5*), fibrosing nonalcoholic steatohepatitis (*6*), and age-related macular degeneration (*7*). Because of its close relationship with wide-ranging diseases, the underlying mechanisms by which cholesterol crystallizes in human bodies have been extensively studied to understand pathogenesis and identify new treatments. It has been traditionally proposed that cholesterol crystals nucleate on the lipid bilayer membranes, such as plasma membranes, lysosomes, and multilamellar bodies, at high cholesterol levels through macrophage models (*3*, *8–10*) and synthetic lipid bilayers (*11–14*). When cholesterol levels exceed the solubility limit in the bilayer membranes, cholesterol forms crystalline cholesterol nanodomains and further grows into triclinic or monoclinic cholesterol monohydrate crystals (*15*, *16*). A triclinic polymorph is thermodynamically stable and has a rhomboidal platy shape, while a monoclinic one is metastable and grows in different shapes, such as a helixes, tubes, rods, or plates (*11*, *17–19*). Cholesterol crystals can nucleate as triclinic or monoclinic polymorphs depending on the kinetic effects (*20*), tissue types (*21*), nucleation sites (*8*), phospholipids (*12*, *13*, *22*), and hydration levels (*23*), but rhomboid-like triclinic crystals predominate over time as the monoclinic form is transformed into the more stable triclinic one.

However, recent 3D visualizations of intact cholesterol crystals inside tissues and cells have revealed that intracellular cholesterol crystals grow in morphologies that differ significantly from their known polymorphic shapes (*24*, *25*). Unlike typical rhomboid-shaped cholesterol crystals, intracellular cholesterol crystals exhibit a thin, sheet-like shape with varying curvatures and rugged edges, forming 3D intertwined agglomerates. In addition, some exhibit a large rod shape and were elongated, extending beyond the size of the cells (*8*, *20*, *25*). Interestingly, these irregularly shaped cholesterol crystals are in contact with lipid droplets, suggesting that unusual crystal morphologies may arise from the unique structural and functional features of the lipid droplets (*20*, *24*). Lipid droplets are central organelles that regulate cellular lipid homeostasis through lipid storage and dynamic interactions with other organelles (reviewed in (*26*)). They have a spherical hydrophobic core containing neutral lipids such as triacylglycerols and cholesteryl esters, covered by a lipid monolayer primarily composed of phospholipids, free cholesterol, and fatty acids (*27*, *28*). In response to cellular cholesterol levels, lipid droplets store free cholesterol on their surfaces and regulate cholesterol concentration through various cellular processes, including intracellular cholesterol trafficking (*29–35*) and enzymatic reactions (*36–39*) (**Fig. 1A**). Therefore, an imbalance in cholesterol homeostasis could lead to excessive cholesterol accumulation and crystallization on the lipid droplet surface. However, cholesterol crystallization on the lipid droplet surface has not been explored extensively, and the mechanisms responsible for the distinctive morphologies of intracellular cholesterol crystals remain a mystery.

**Fig. 1.**
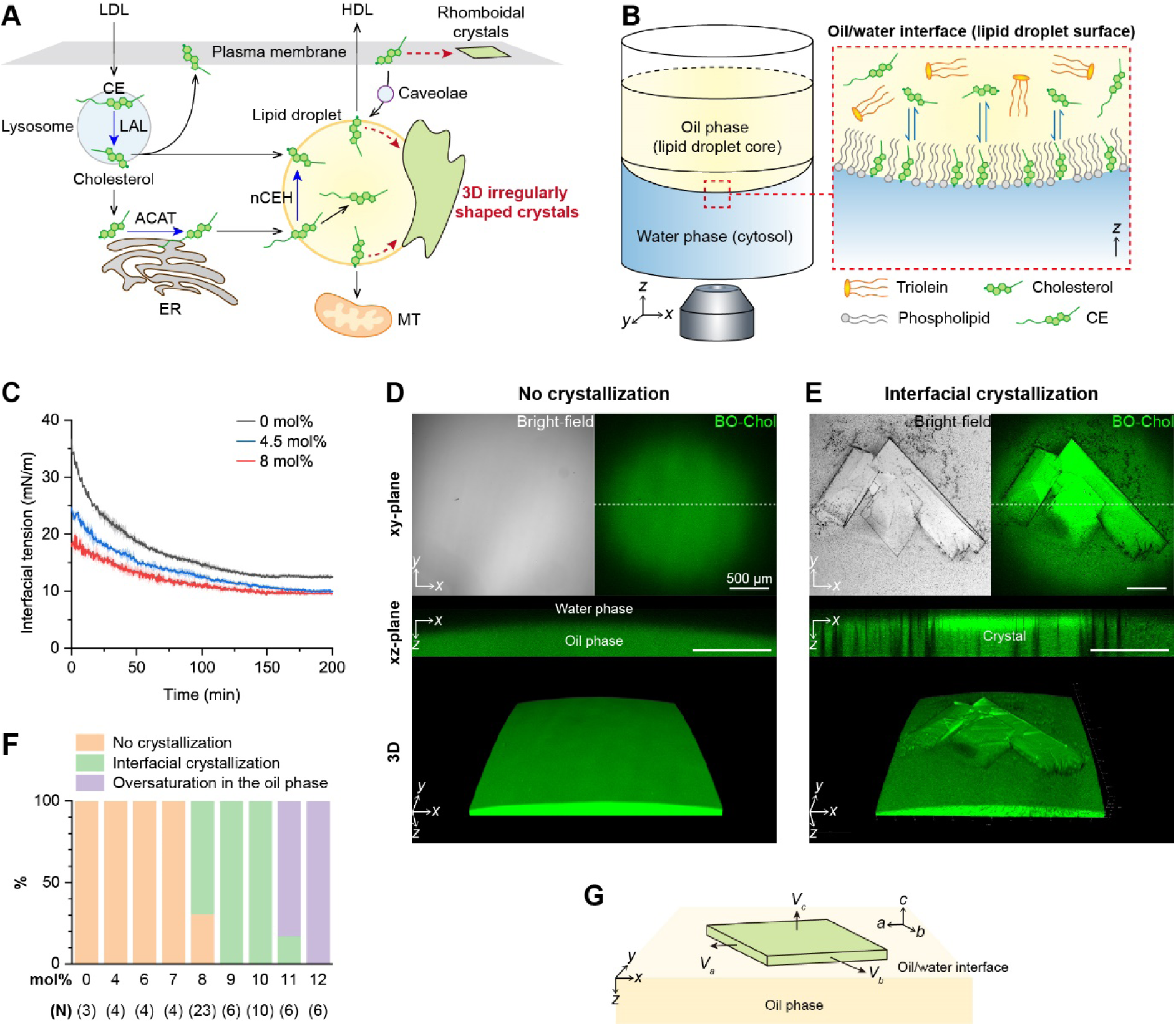
Cholesterol crystallization on the model lipid droplet surface. **(A)** Intracellular cholesterol processing. Cholesteryl esters (CEs) enter the cell via low-density lipoproteins (LDLs) and are hydrolyzed into free cholesterol by lysosomal acid lipase (LAL) in the lysosome. Free cholesterol moves to the endoplasmic reticulum (ER), re-esterified into CE by acyl-coenzyme A:cholesterol acyltransferase (ACAT), and stored in lipid droplets. Cholesterol levels in lipid droplets are regulated by converting CEs back to cholesterol with neutral cholesterol ester hydrolase (nCEH), cholesterol efflux to high-density lipoproteins (HDLs), and cholesterol transport among lipid droplets, plasma membrane, lysosomes, and mitochondria. Excess cholesterol forms rhomboidal platy crystals on the plasma membrane and three-dimensionally irregular crystals on lipid droplets. **(B)** Schematic illustration of the lipid droplet model for cholesterol crystallization visualization. **(C)** The variation in interfacial tension at different cholesterol concentrations of the oil phase (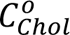). Standard deviation is indicated by shaded regions. **(D-E)** Oil/water interfaces without **(D)** and with cholesterol crystals **(E)** were examined using bright-field and BO-Chol fluorescence imaging. The xz-plane views represent the 2D cross-sections at the dotted lines of the xy-plane images. **(F)** Probability of interfacial cholesterol crystallization at different 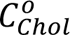 based on multiple experimental trials (N). (**G**) Scheme of growth of a plate-like cholesterol crystal at the oil/water interface.

Here, we reveal how cholesterol crystallizes into non-rhomboidal crystals on the surface of lipid droplets and eventually evolves into a 3D irregular sheet-like architecture to understand the non-traditional morphological growth of cholesterol crystals observed adjacent to lipid droplets. Our findings highlight that the interfacial transport kinetics, unique lipid composition, and curved structure of the lipid droplets significantly impact on interfacial cholesterol crystallization, resulting in the coalescence of non-rhomboidal platy crystals into 3D sheet-like agglomerates.

## Results

### Nucleation and growth of platy crystals at the interface

To monitor cholesterol crystallization on the lipid droplet surface in real time through confocal fluorescence microscopy with 3D image processing, we established an *in vitro* reconstitution system that mimics the lipid droplet and its surrounding environment (**Fig. 1B**). This model system comprises three components: nonpolar oil phase, water phase, and oil/water interface. The oil phase mimics the lipid droplet core by containing triolein, cholesterol, cholesteryl esters, and phospholipids. Traces of fluorescently labeled cholesterol (BO-Chol) and phospholipid (TR-DHPE) were also included in the oil phase as fluorescence probes for cholesterol and phospholipids. The water phase is a physiologically relevant aqueous buffer solution that contains major salts present in cytosol at pH 7.4. We formed this model in the 96-well cell culture plate by simply placing the water phase on the bottom and then gently dropping the oil phase on top of the water phase, which led to the formation of the oil/water interface that mimics the lipid droplet surface.

We first considered the basic system where the oil phase only consisted of triolein and cholesterol. Because of its amphiphilic nature, cholesterol conceivably diffuses from the oil phase to the oil/water interface, forming a lipid monolayer with its hydroxyl group oriented toward the water phase. Based on Fick’s first law of diffusion, increasing the cholesterol concentration of the oil phase 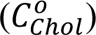 increases its concentration gradient along the direction of oil-to-interface diffusion, potentially leading to an increase in the interfacial cholesterol density 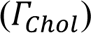. Consistent with this expectation, Interfacial tension measurement showed that as 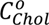 increased, the interfacial tension at the oil/water interface decreased more rapidly, reaching as low as 10 mN/m at equilibrium (**Fig. 1C**), which indicates a larger and faster accumulation of cholesterol at the interface with the increase in of 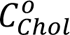.

To determine if cholesterol that accumulates at the oil/water interface crystallizes, we examined the oil/water interface while increasing 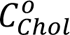 from 0 mol% to the solubility limit of approximately 12 mol% (**Fig. 1D, 1E, 1F, and S1A**). The two-dimensional (2D) and three-dimensional (3D) BO-Chol fluorescence images clearly identified the location of the curved oil/water interface due to a significant difference in the BO-Chol fluorescence intensity between the oil and water phases (**Fig. 1D**). We did not find any crystallization at 7 mol% and below for 14 hours (**Fig. 1F**) but observed that at 8 mol%, angular plate-like crystals began to appear at the oil/water interface with a high probability, forming a clear threshold (**Fig. 1E**). The straight crystal edges were clearly distinguished through bright-field microscopy because of the different refractive indexes between the crystals and surrounding media. Cholesterol crystals were also detected in fluorescence imaging, with most of them exhibiting a higher BO-Chol fluorescence intensity than the surrounding media. The cross-sectional image (xz-plane) showed that the crystals were located at the oil/water interface and had a platy shape parallel to the interface with a width/thickness ratio of more than 10. This indicates that crystals grow faster parallel to the interface (along the a- and b-axes) than vertically (along the c-axis) (**Fig. 1G**).

When 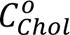 exceeded 8 mol%, the interfacial cholesterol crystals grew larger than the field of view (**Fig. S1A**). When 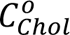 reached 11 mol% or higher, the oil phase became oversaturated with cholesterol, and needle-like crystals formed in the oil phase (**Fig. S1A and S1B**). These results indicate that cholesterol crystallizes first at the surface of the lipid droplet rather than in the core, forming plate-like crystals oriented parallel to the surface.

### Crystal elongation depending on interfacial cholesterol diffusion rate

To characterize the overall shape of cholesterol crystals, we measured the size and angle of interfacial cholesterol crystals formed at 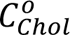 of 8 mol% (**Fig. 2A, 2C, and 2D**). We observed that fewer than ten cholesterol crystals nucleated at the interface, growing gradually larger in different shapes and rates (**Fig. S2A and S2B**). The size quantification showed that the cholesterol crystals tended to elongate with various edge ratios between 2 and 20, and some of them reaching a few millimeters in edge length (**Fig. 2C**). This elongated morphology differs from the rhomboid-like shape of triclinic cholesterol monohydrate crystals formed in atherosclerotic lesions (*8*, *9*, *21*) and model lipid bilayers (*11*, *13*). The morphological difference can result from the polymorphism of cholesterol crystals, which form five different crystallographic structures that exhibit different anisotropic growth (*40*). However, based on the XRD data, elongated crystals are also triclinic polymorphs, the same as the rhomboidal ones. The XRD profile of the elongated crystals displayed the same prominent peaks corresponding to the D-spacing of 34.2 Å and 17.1 Å (**Fig. S3**), aligning with the diffraction pattern of triclinic monohydrate polymorph (*41*). In addition, they have the same specific bimodal angles of ∼80° (α) and ∼100° (β) (**Fig. 2D**), consistent with triclinic cholesterol monohydrate crystals with the largest angle of a unit cell being 100.8° (*42*, *43*).

**Fig. 2.**
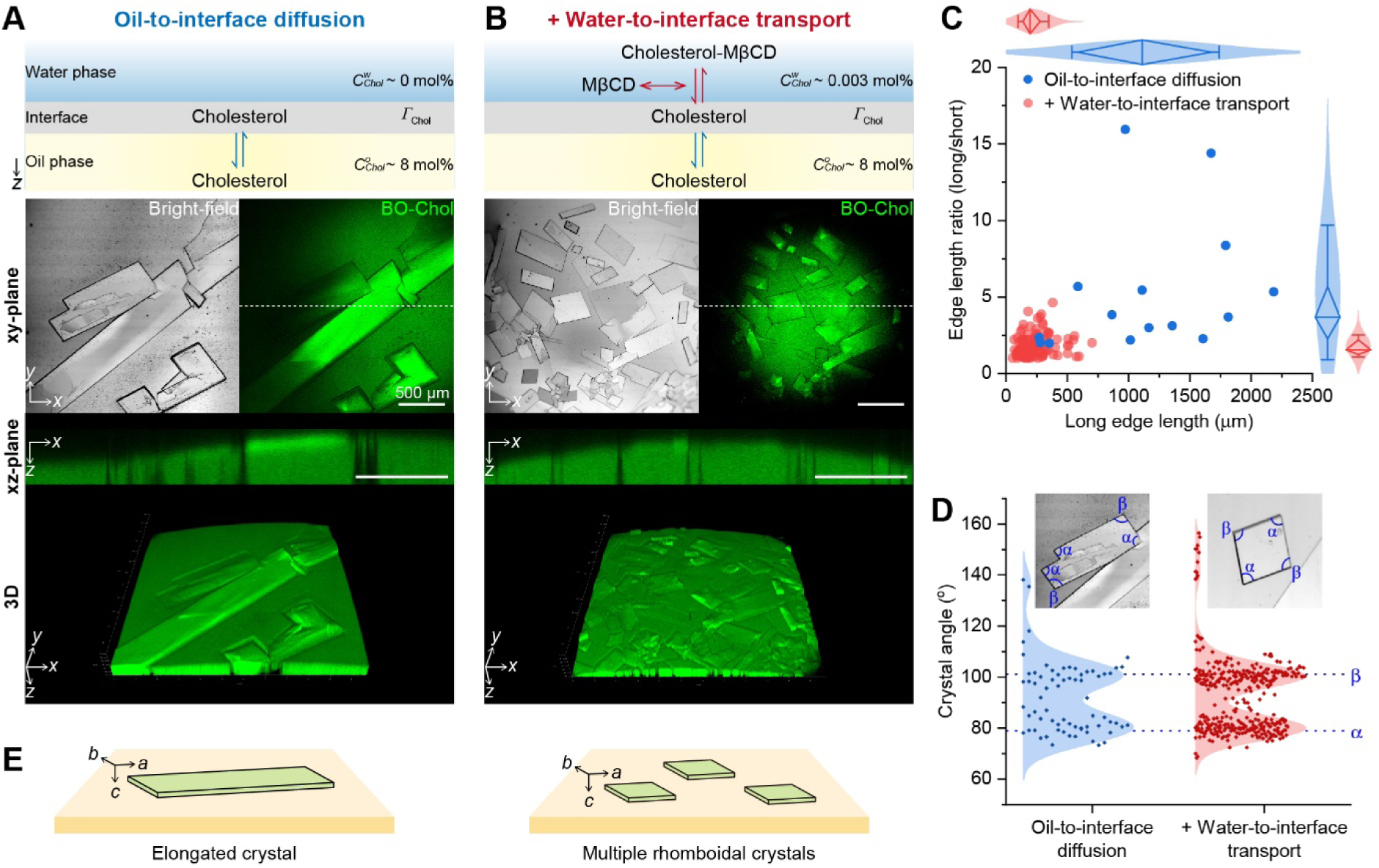
Effect of diffusion kinetics on interfacial cholesterol crystallization. **(A)** Interfacial cholesterol crystals were observed using bright-field and BO-Chol fluorescence imaging at 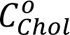 of 8 mol%. Cholesterol accumulated at the oil/water interface exclusively via only free diffusion from the oil phase to the interface. The xz-plane views represent the 2D cross-sections at the dotted lines of the xy-plane images. (**B**) Cholesterol was additionally transferred to the interface via MβCD-mediated water-to-interface transport. **(C)** The edge length of cholesterol crystals was displayed as a scatter plot. The length ratio of long and short sides was plotted against the long edge length. The distribution of these measurements was approximated using a normal distribution. Results are displayed using diamond box plots, which include a median line and whiskers representing the standard deviation. **(D)** The distribution of angles within cholesterol crystals was estimated using the Kernel distribution method. Peaks α and β of the angle distribution were identified at 79° and 101°, respectively, determined through second derivatives of the distribution. The samples analyzed in **C** and **D** represent cholesterol crystals that had grown for 12-14 hours following the formation of the oil/water interface. **(E)** The schematic illustration of elongated and rhomboidal crystals growing along the a-, b-, and c-axes at the oil/water interface. Three axes (a, b, and c) parallel different crystal edges.

Another possibility is diffusion-limited crystal growth kinetics. Cholesterol crystals can grow at the oil/water interface through a series of steps: cholesterol molecules diffuse from the oil phase to the oil/water interface, move to the crystal surface, are stripped from the surrounding solvent molecules, move along the crystal surface to a suitable lattice position, and finally integrate into the crystal structure (*44*). If the diffusion of cholesterol molecules from the oil phase to the crystal surface located at the oil/water interface is slower than their integration, the concentration gradient along the crystal surface can be generated, potentially leading to the different crystal growth rates along the a- and b-axes and the elongated morphology (**Fig. 2E**). On the other hand, when diffusion is faster than integration, cholesterol molecules can be evenly distributed on the crystal surface, resulting in comparable growth rates along the a- and b-axes and a rhomboidal crystal shape.

In order to test whether the diffusion rate affects the crystal shape, we increased the transport of cholesterol to the oil/water interface by using MβCD, which encapsulates cholesterol in the water phase and reversibly transfers it between the water phase and the oil/water interface (*45*) (**Fig. 2B and S2C**). We saturated the MβCD-including water phase with cholesterol to enhance the diffusion of cholesterol molecules into the oil/water interface without changing 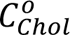. We confirmed that the MβCD-mediated transport increased the adsorption rate and density of cholesterol at the oil/water interface, as indicated by the faster reduction of the interfacial tension to ∼9 mN/m around 200 minutes (**Fig. S4**). Our imaging showed that the MβCD-mediated cholesterol transport induced dozens of crystal nucleation events at the interface (**Fig. 2B**), and the nucleated crystals grew in a rhomboid-like shape with an edge ratio of approximately 2 on average (**Fig. 2D**) and consistent bimodal angles (**Fig. 2E**). Based on the XRD data (**Fig. S3B**), these rhomboidal crystals were also proven to be a triclinic cholesterol monohydrate polymorph. These findings suggest that triclinic cholesterol crystals grow in a rhomboidal shape under rapid diffusion of cholesterol to the oil/water interface, whereas they develop an elongated morphology under slow diffusion. The elongation of the cholesterol crystals may be the possible mechanism behind the formation of rod-like crystals that could pierce cell membranes (*8*, *24*, *25*).

### Variation in crystal angle with cholesteryl esters

We next addressed the question of whether interfacial cholesterol crystallization is impacted by cholesteryl esters, which are major components of lipid droplets. Cholesteryl esters comprise 10-50% of lipid droplets but are not present in the plasma membranes (*46–48*), potentially explaining the differences in cholesterol crystallization between these locations. We first included cholesteryl palmitate (CP), which is one of the representative saturated cholesteryl esters, in the oil phase. CP mixed well with triolein at CP concentration in the oil phase 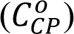 below 5 mol% but formed platy crystals at the oil/water interface at 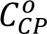 of 5 mol% or more (**Fig. 3A** and **S5**). CP crystals were formed parallel to the interface as well as at angles not parallel to the interface (triangles in **Fig. 3A**). They had non-uniform shapes with specific three angles of 54° (γ), 103° (δ), or 126° (ε) (**Fig. 3B**) and three main Bragg peaks corresponding to D-spacings of 52.65 Å, 26.52 Å, and 17.70 Å (**Fig. 3C and Table S1**), differing from the Bragg peaks of cholesterol crystals.

**Fig. 3.**
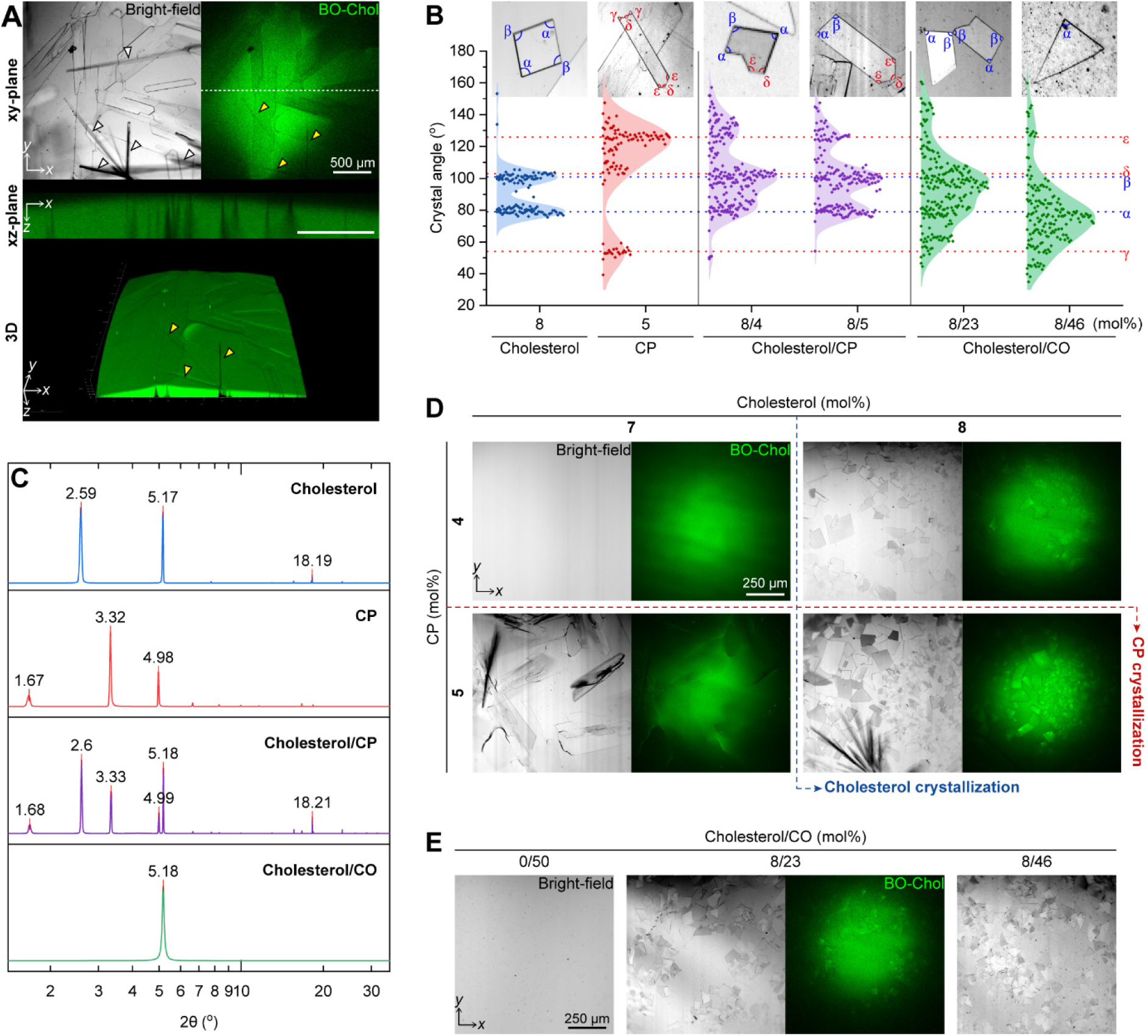
Interfacial cholesterol crystallization with cholesteryl esters. **(A)** Interfacial CP crystallization was observed using bright-field and BO-Chol fluorescence imaging. The xz-plane images represent cross-sections at the dotted lines of the xy-plane images. The white and yellow triangles indicate the CP crystals growing non-parallel to the oil/water interface and their contact sites at the interface, respectively. **(B)** The angle distributions of cholesterol and CP crystals were analyzed using the Kernel distribution. Peaks α and β, corresponding to 79° and 101°, are the angle distribution peaks for cholesterol crystals. Peaks γ, δ, and ε, at 54°, 103°, and 126°, respectively, represent the angle distribution of CP crystals. These peak values were derived from second derivatives of the distribution. The analyzed samples are crystals that had grown for 12-14 hours following the formation of the oil/water interface. **(C)** X-ray diffraction analysis of cholesterol crystals formed in the presence of cholesteryl esters. **(D-E)** Representative bright-field and fluorescence images of interfacial crystals formed with different concentrations of cholesterol and cholesteryl esters, including CP (**D**) and CO (**E**). The oil phase contained cholesterol and cholesteryl esters, while the water phase contained MβCD.

By controlling both cholesterol and CP concentrations in the oil phase, we confirmed that CP had minimal impact on the critical cholesterol concentration for its interfacial crystallization; cholesterol crystals consistently appeared at 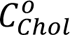 of 8 mol% or more, independent of CP concentration (**Fig. 3D**). However, in the presence of CP, a significant increase in cholesterol crystal nucleation was observed at the oil/water interface, indicating the potential role of CP as a nucleating agent for cholesterol crystals, which is consistent with the previous computational approach (*20*). Furthermore, it is notable that cholesterol crystals primarily displayed an irregular, non-quadrilateral shape, characterized by three dominant angles of around 79°, 101°, and 126° (**Fig. 3B and 3D**). The difference in the shape of cholesterol crystals with and without CP suggests that cholesterol may co-crystallize with CP within the same crystal structure. The XRD data corroborate this possibility, showing the Bragg peaks of both triclinic cholesterol monohydrate and CP crystals within the mixed cholesterol crystals formed with CP (**Fig. 3C and Table S1**). This integrated cholesterol/CP structure is expected to exhibit the physical characteristics of both components, resulting in a combination of angles characteristic of cholesterol crystals (79° and 101°) and CP crystals (103° and 126°).

We also tested another representative cholesteryl ester, cholesteryl oleate (CO). CO is an unsaturated cholesteryl ester and does not crystallize in the oil phase even at a high concentration of 50 mol% at physiological temperature (**Fig. 3E**). Similar to CP, CO promoted cholesterol crystallization, as evidenced by the numerous cholesterol crystals forming at 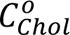 of 8 mol%. However, unlike the sharp angle distribution of cholesterol/CP crystals, the cholesterol crystals formed with CO exhibited a broad spectrum of angles (**Fig. 3B**). As the CO concentration increased, the angular distribution shifted lower, centering around 79°, yet remained wide-ranging. Despite their irregular shape, the crystals still had a Bragg peak for triclinic cholesterol monohydrate crystals, corresponding to the D-spacing of 17.1 Å (**Fig. 3C and Table S1**). Our experiments demonstrate that cholesteryl esters, including CP and CO, actively influence both the nucleation and the resulting shape of cholesterol crystals, potentially leading to the formation of irregularly shaped triclinic cholesterol monohydrate crystals on the surface of lipid droplets.

### Phospholipid-mediated recruitment of cholesterol to the interface

The variation in cholesterol crystallization by cholesteryl esters prompted us to ask whether cholesterol crystallization is also affected by phospholipids that cover the lipid droplet surface. Phospholipids were observed to accumulate at the oil/water interface, as indicated by the high TR-DHPE fluorescence intensity at the oil/water interface (**Fig. 4A**). BO-Chol fluorescence images illustrated the colocalization of cholesterol with phospholipids at the interface, suggesting the recruitment of cholesterol to the phospholipid-laden interface through intermolecular interactions, including hydrogen bonds between the head group of phospholipids and the hydroxyl group of cholesterol (*49*). The interfacial tension measurement showed that phospholipids and cholesterol rapidly lowered interfacial tension to approximately 1 mN/m (**Fig. S6**), aligning with the reported surface tension of the lipid droplets (approximately 1-2 mN/m) (*50*).

**Fig. 4.**
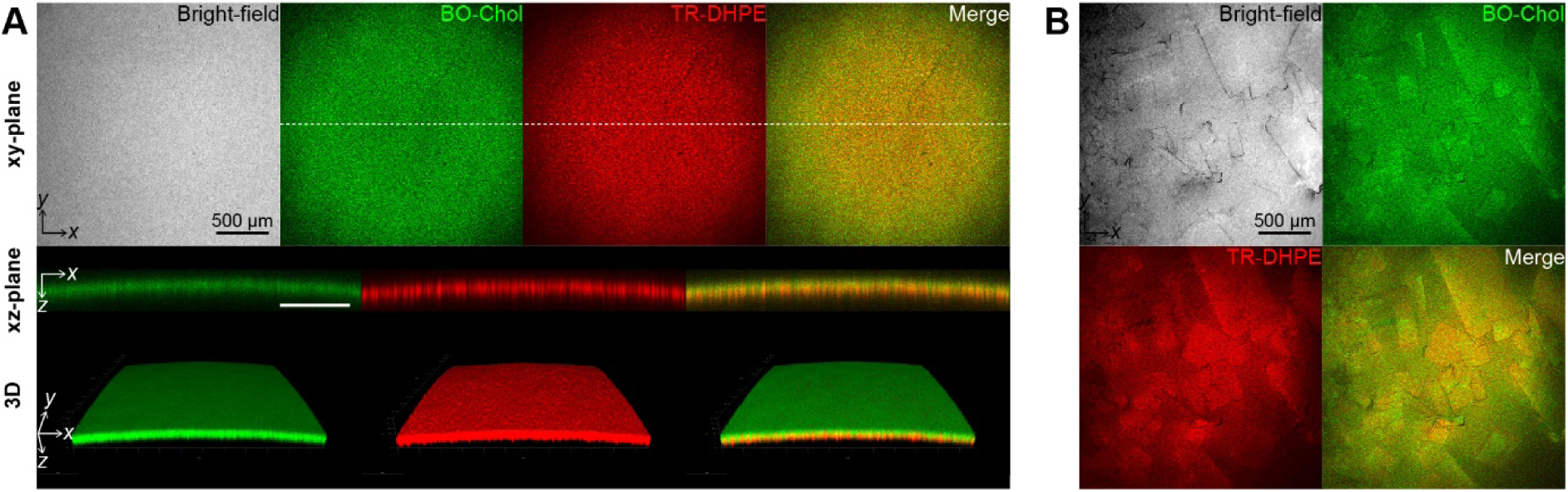
Cholesterol crystallization at the phospholipid-laden oil/water interface. The oil/water interface was observed using bright-field and fluorescence imaging in the presence of cholesterol and phospholipids. The concentrations of cholesterol/phospholipids in the oil phase were 8/2 mol% (**A**) and 10/2 mol% (**B**). The xz-plane images represent the cross-section at the dotted line of the xy-plane images.

By controlling the cholesterol concentration, we identified that unlike phospholipid-free interface, cholesterol did not crystallize at the phospholipid-laden interface at 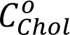 of 8 mol% (**Fig. 4A**) and that a higher 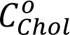 (10 mol% and more) is required for crystallization (**Fig. 4B**). The cholesterol crystals grew in a quadrilateral shape but had partially rounded angles (**Fig. 4B and S7)**. Initially, the TR-DHPE fluorescence intensity of the cholesterol crystals was lower than that of the phospholipid-laden interface, but it gradually increased and eventually surpassed the interface (**Fig**. **S7**). This enrichment of phospholipids on the crystal surface represents their preferential interaction with cholesterol crystals, which is consistent with previous observations of phospholipid-covered triclinic monohydrate crystals (*13*). Thus, our results suggest that the preferential interaction of phospholipids with cholesterol boosts the interfacial density of cholesterol but inhibits interfacial crystallization and affects the shaping of cholesterol crystals.

### Macroscopic crystal growing to sheet morphology

The cholesterol crystals observed so far are over ten times smaller than our model lipid droplets, but cholesterol crystals can grow to scales larger than lipid droplets in a cell, potentially forming macroscopic structures that could pierce cell membranes (*8*, *24*, *25*). To explore the macroscopic growth process of cholesterol crystals, we developed cholesterol crystals at scales comparable to our model lipid droplets by increasing 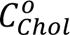 to more than 8 mol% and observed their 3D structure at the oil/water interface. Our visualization showed that interfacial cholesterol crystals can macroscopically develop into flat or curved sheet-like crystals, depending on the nucleation frequency at the oil/water interface.

When a few cholesterol crystals occurred at the oil/water interface without MβCD, the crystals grew in a flat platy shape, unaffected by the positive Gaussian curvature of the interface (**Fig. 5A**). The cholesterol crystals flattened the curved oil/water interface upon contact and formed angular interfaces by meeting at non-parallel angles (yellow triangle in **Fig. 5A**). Although the edge of cholesterol crystals tended to roughen in the presence of phospholipids, cholesterol crystals still expanded two-dimensionally into sheet-shaped forms, altering the adjacent interface curvature from concave to convex along the z-axis (**Fig. 5B and S8**). As 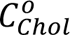 increased, cholesterol crystals expanded to cover most of the interface area (**Fig. 5C and S9**). When the cholesterol crystals outgrew the limited area of the interface, they were folded and crumpled, resulting in a jagged structure of both cholesterol crystals and interface (**Fig. 5C**). This dynamic deformation of the oil/water interface, observed during the macroscopic sheet-like crystal growth, is possibly attributed to the extremely low interfacial energy at the phospholipid-laden interface (**Fig. 4B**). These results suggest that cholesterol crystals can grow in massive, sheet-like morphology at the lipid droplet surface, distorting the curvature of the lipid droplets and escaping from the lipid droplets (**Fig. 5D**).

**Fig. 5.**
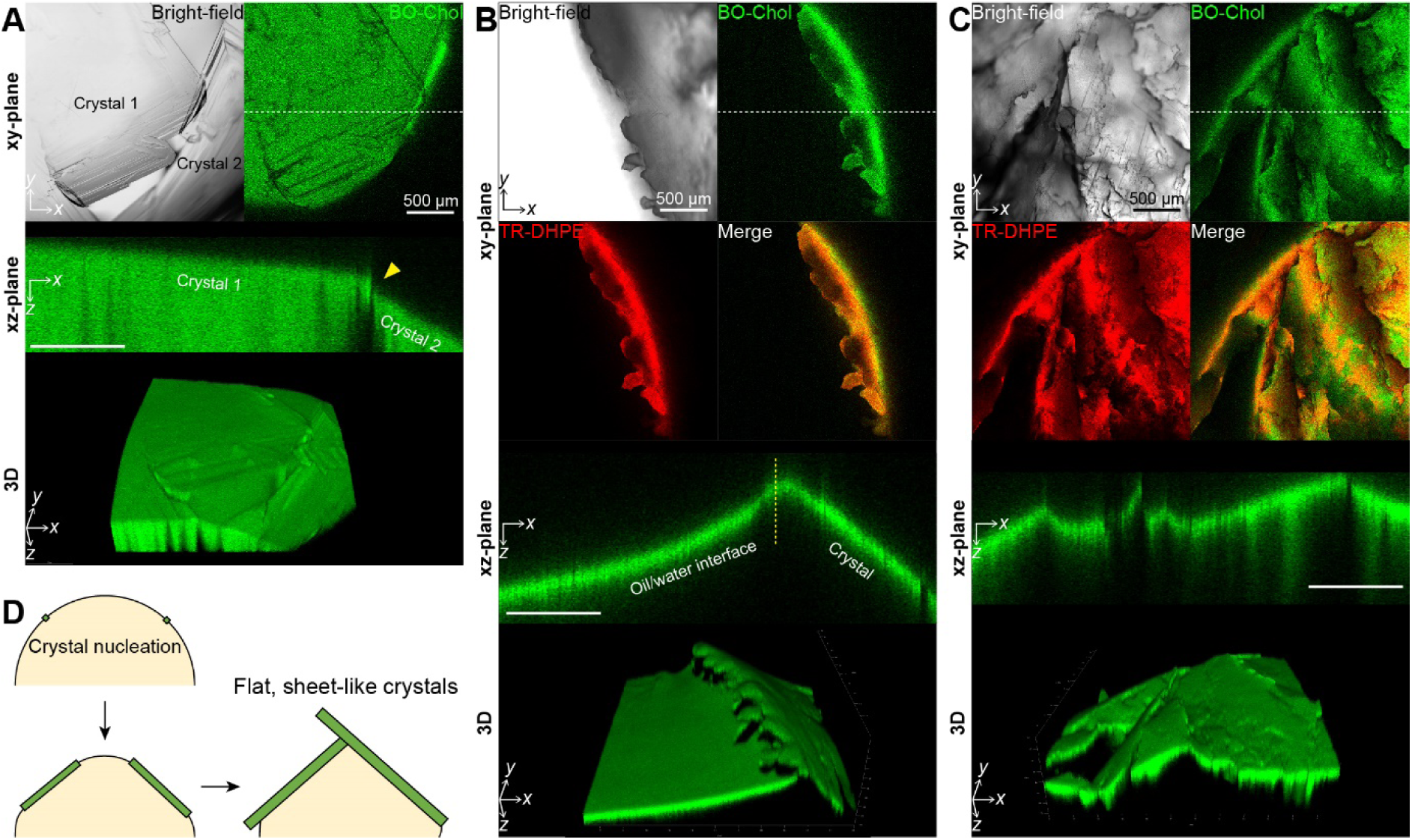
Flat, sheet-like cholesterol crystals grown at the oil/water interface. **(A-C)** The macroscopically grown cholesterol crystals at the oil/water interface were visualized by the bright-field and fluorescence imaging at cholesterol/phospholipid concentrations of 9/0 mol% **(A)**, 12/2 mol% **(B)**, and 14/2 mol% **(C)**. The xz-plane images represent the cross-section at the dotted line of the xy-plane image. **(D)** Schematic illustration of the presumable nucleation-and-growth process of cholesterol crystals on the lipid droplet surface.

### Development of 3D curved crystal agglomerates

However, when many crystals nucleate and grow simultaneously on the confined lipid droplet surface, they are likely to meet and influence the growth process of each other. Real-time visualization of directly contacting cholesterol crystals revealed that different crystals readily merged upon contact, maintaining their orientation without undergoing 2D rotational movements (**Fig. 6A**). Upon merging, the crystals expanded two-dimensionally, filling the interface near the contact point and thereby extending the length of the contact edge. When the crystals partially overlapped, they grew by enlarging the area of overlap (**Fig. 6B**). The crystal overlaps sometimes resulted in the dissolution of one of the crystals (**Fig. 6C**). These results suggest that cholesterol crystals readily combine in a random orientation to form an irregular crystal aggregate, thereby losing original shapes (**Fig. 6D**). As 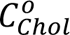 increased, the number of nucleated cholesterol crystals increased, leading to their rapid, extensive coalescence at the oil/water interface and the creation of a large, curved, sheet-like aggregate (**Fig. S10**). These results suggest that the lateral aggregation of cholesterol crystals on the curved lipid droplet surface promotes the formation of three-dimensionally curved sheet-like crystals (**Fig. 6E**), while an independent cholesterol crystal grows in flat sheet-like forms.

**Fig. 6.**
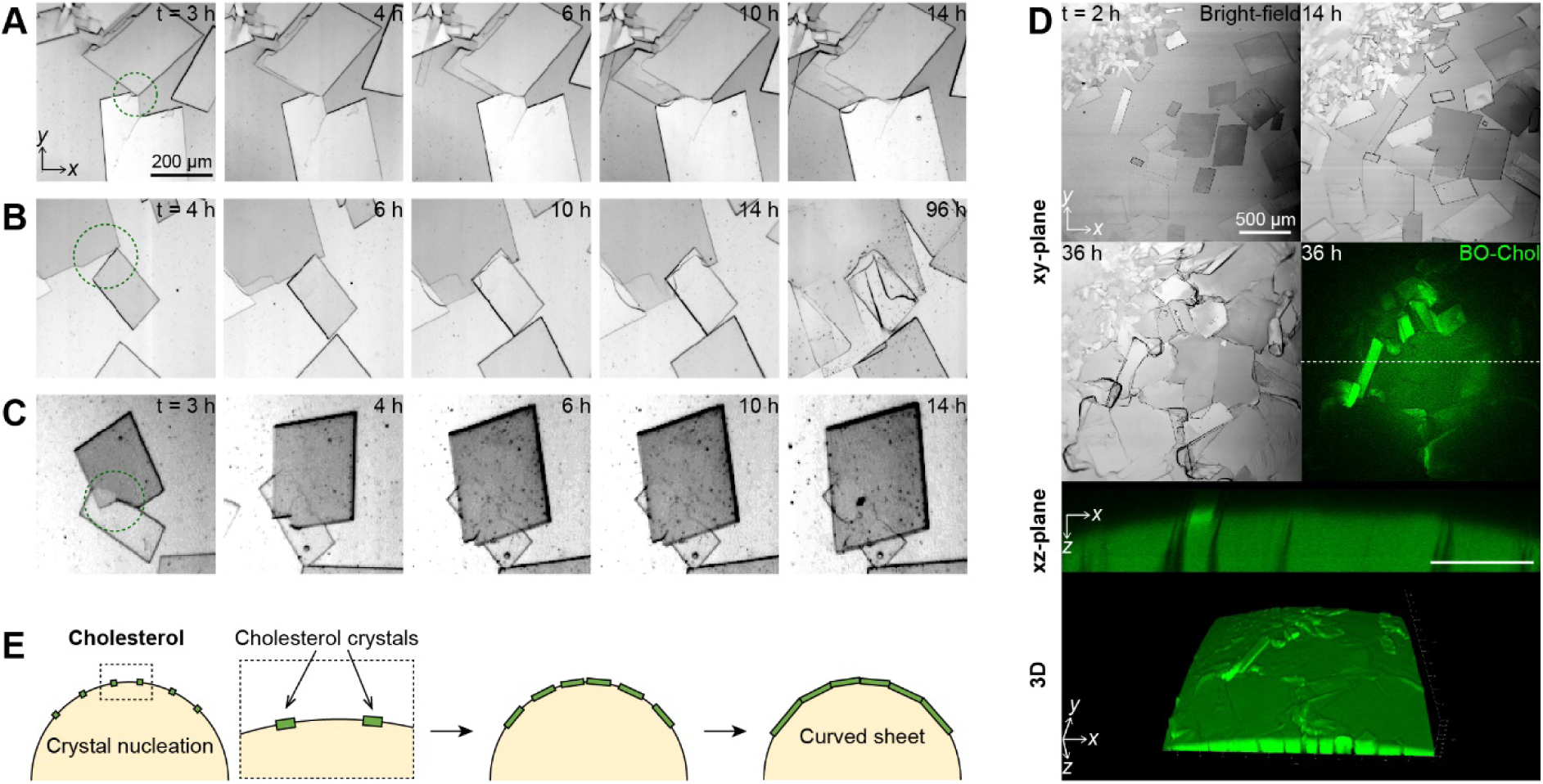
Aggregation of cholesterol crystals into a curved, sheet-like agglomerate at the oil/water interface. **(A-C)** Cholesterol crystals meet each other at the oil/water interface through lateral merging (**A**), partial overlap (**B**), and dissolution (**C**). **(D)** The aggregation of cholesterol crystals was visualized by the bright-field and fluorescence imaging at a 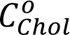 of 9 mol% with the MβCD-mediated water-to-interface transport. **(E)** Schematic illustration of the aggregation of cholesterol crystals on the curved oil/water interface.

Considering the extremely low surface energy (γ ∼1 mN/m) and the limited area of the lipid droplet surface, sheet-like cholesterol crystals may escape from the lipid droplets and merge with those originating from different lipid droplets in the confined intracellular space. This process results in the formation of 3D entangled cholesterol crystal architectures, as observed in previous studies (*24*, *25*).

## Discussion

This study demonstrates the unique morphological growth processes of cholesterol crystals on the surface of lipid droplets. Our results propose that the lipid droplet surface is an active site in which cholesterol crystallizes into non-rhomboidal plate-like crystals and even 3D sheet-like agglomerates. This provides fundamental insights into understanding the distinctive morphology of intracellular cholesterol crystals observed in recent studies (*8*, *20*, *24*, *25*). Using an *in vitro* model lipid droplet system (**Fig. 1**), we highlight that these crystals can grow in different morphologies from the rhomboidal cholesterol crystals originating from the plasma membranes, depending on the unique features of the lipid droplet surface: 1) interfacial cholesterol transport (**Fig. 2**), 2) lipid composition (**Fig. 3 and 4**), and 3) the spherical structure of the lipid droplets (**Fig. 5 and 6**).

Active interactions between lipid droplets and other organelles may also affect cholesterol crystallization. Direct membrane contacts of lipid droplets with the lysosome, endoplasmic reticulum, and mitochondria can cause interfacial lipid transfers on the lipid droplet surface (*34*, *35*), potentially inducing the formation or dissolution of cholesterol crystals. Moreover, the hydrolysis of cholesteryl esters to free cholesterol via neutral cholesterol ester hydrolases (*36–38*) and lysosomal acid lipase (*39*, *51*) could influence the cholesterol crystallization by changing the cholesterol accumulation rate and the lipid composition at the lipid droplet surface. Therefore, further studies on the role of other cellular organelles and enzymatic reactions in regulating cholesterol crystallization could provide a more extensive picture of the physiological and pathological contexts.

In summary, this study elucidates the 3D morphological growth dynamics of cholesterol crystals nucleating from the surface of lipid droplets. Real-time imaging with a meticulously controlled lipid droplet model system has uncovered significant influences of interfacial transport, lipid composition, and the geometry of the lipid droplet surface on the crystallization and shaping of cholesterol crystals. Our results provide novel insights into the fundamental mechanisms of the formation and non-traditional growth of cholesterol crystals, distinct from those derived from plasma membrane.

## Materials and Methods

### Materials

Triolein, cholesterol, cholesteryl palmitate, and cholesteryl oleate were purchased from Sigma-Aldrich. Phospholipids including 1,2-dioleoyl-sn-glycero-3-phosphocholine (DOPC), 1,2-dioleoyl-sn-glycero-3-phosphoethanolamine (DOPE), and 23-(dipyrrometheneboron difluoride)-24-norcholesterol (BO-Chol) were purchased from Avanti Polar Lipids. The fluorescent phospholipid probe, Texas Red 1,2-dihexadecanoyl-sn-glycero-3-phosphoethanolamine (TR-DHPE), was purchased from Thermo Fisher Scientific. Components for the aqueous phase, including sodium chloride, potassium chloride, magnesium chloride, 4-(2-Hydroxyethyl)piperazine-1-ethanesulfonic acid (HEPES), and methyl-beta-cyclodextrin (MβCD), were purchased from Sigma-Aldrich. Experiments were conducted in transparent 96-well cell culture plates purchased from SPL Life Sciences.

### Preparation of lipid droplet model system and visualization

The oil phase mimics the lipid droplet core, consisting of triolein (46-100 mol%), cholesteryl esters (0-46 mol%), cholesterol (0-14 mol%), and phospholipids (0-2 mol%), where phospholipids consisted of 1.5 mol% DOPC and 0.5 mol% DOPE. To monitor the distributions of cholesterol and phospholipids, BO-Chol and TR-DHPE were added to the oil phase at a concentration of 2 × 10^-4^ mol% and 5 × 10^-3^ mol%, respectively. The lipid-containing oil mixture was incubated at 50°C for 1 hour to ensure complete dissolution of the lipids, and then cooled down to 37°C. The cytosol-mimicking aqueous phase included 12 mM sodium chloride, 145 mM potassium chloride, 2 mM magnesium chloride, and 10 mM HEPES, adjusted to pH 7.4.

In order to induce interfacial cholesterol transport between the water phase and interface, cholesterol and MβCD were added to the water phase at concentrations of 0.15 mM and 3 mM (1:20 mol%), respectively. At this concentration condition, cholesterol was almost saturated in the water phase. The MβCD-mediated cholesterol transport barely changed the cholesterol concentration in the bulk oil phase (8 mol%) because the amount of cholesterol encapsulated by MβCD in the water phase is extremely low (approximately 0.003 mol%).

For the experimental setup and imaging, the transparent 96-well cell culture plate was placed on the stage of confocal microscope (STELLARIS 5, Leica) and heated to 37°C with the Stage-top Incubator System T (Live Cell Instrument). Initially, 100 µl of the aqueous phase was injected on bottom of a well of the cell culture plate and allowed to equilibrate to 37°C. Then, 400 µl of the oil phase was gently deposited on the water phase to establish the oil/water interface, marking the onset of the experiment (t = 0). Cholesterol crystals formed at the oil/water interface were typically visualized for 14 hours through combined bright-field and fluorescence imaging modes. 3D images were acquired by a z-stack scan mode. The crystal sizes and angles in the obtained images were analyzed by ImageJ software (*52*).

### Pendant drop analysis

Interfacial tension measurements were performed using the pendant drop method. Using the SEO Phoenix 300 Touch equipment, droplets from the water phase were suspended in the lipid oil phase composition described previously. Images of the droplet were captured at intervals of several seconds using the equipment, continuing until the droplet shape stabilized. The recorded images were then analyzed to determine the interfacial tension. Calculations were conducted using Surfaceware 9 software, supplemented with custom scripts written in MATLAB. To ensure the reproducibility of the results, the entire sample preparation and measurement procedures were repeated at least three times.

### X-ray diffraction (XRD)

XRD analysis was performed using a Rigaku Smartlab High-resolution Powder X-Ray Diffractometer with a Cu Kα radiation source (λ=1.5406 Å). Theta-theta measurements spanned a 2θ range from 1° to 80°, employing a step size of 0.01° and a scan rate of 10° per minute. The diffractometer was operated at 45 kV and 200 mA, under room temperature conditions. For the sample preparation, crystals formed at the oil/water interface were carefully transferred onto filter paper and isolated by removing the water and oil using a vacuum freeze dryer. The remaining crystal samples on the filter paper were then mounted on zero-background silicon sample holders designed for XRD analysis. Following data collection, XRD patterns were analyzed using Rigaku XRD PDXL software. The crystal structures were identified by matching the observed diffraction peaks with the obtained Rigaku RAW data. Both first and second derivative analyses of the peaks were also checked to find hidden peaks. To ensure reproducibility of the results, the entire sample preparation and measurement procedures were repeated at least three times.

## Acknowledgements

This research was supported by the KAIST Institute of Technology Value Creation, Industry Liaison Center (G-CORE Project) grant funded by the Ministry of Science and ICT (MSIT) (N11210011), the Basic Science Research Program through the National Research Foundation of Korea (NRF) funded by the Ministry of Education (2021R1A6A3A01086522), and NRF grant funded by the MSIT (RS-2023-00276535).

## Author contributions

HRL and SK contributed equally to this work. HRL, SK, and SQC conceived the idea and designed the research. HRL and SK performed the experiments and analyzed data. HRL, SK, and SQC wrote the manuscript.

## Data and materials availability

The authors declare that the main text and Supplementary Materials files contain all relevant data. Raw data are available on request from the corresponding author.

