## Supplementary Materials for "Lipid droplet surface promotes three-dimensional morphological evolution of non-rhomboidal cholesterol crystals"

Hyun-Ro Lee *et al.*

**This PDF file includes:**

Figs. S1 to S10
Tables S1

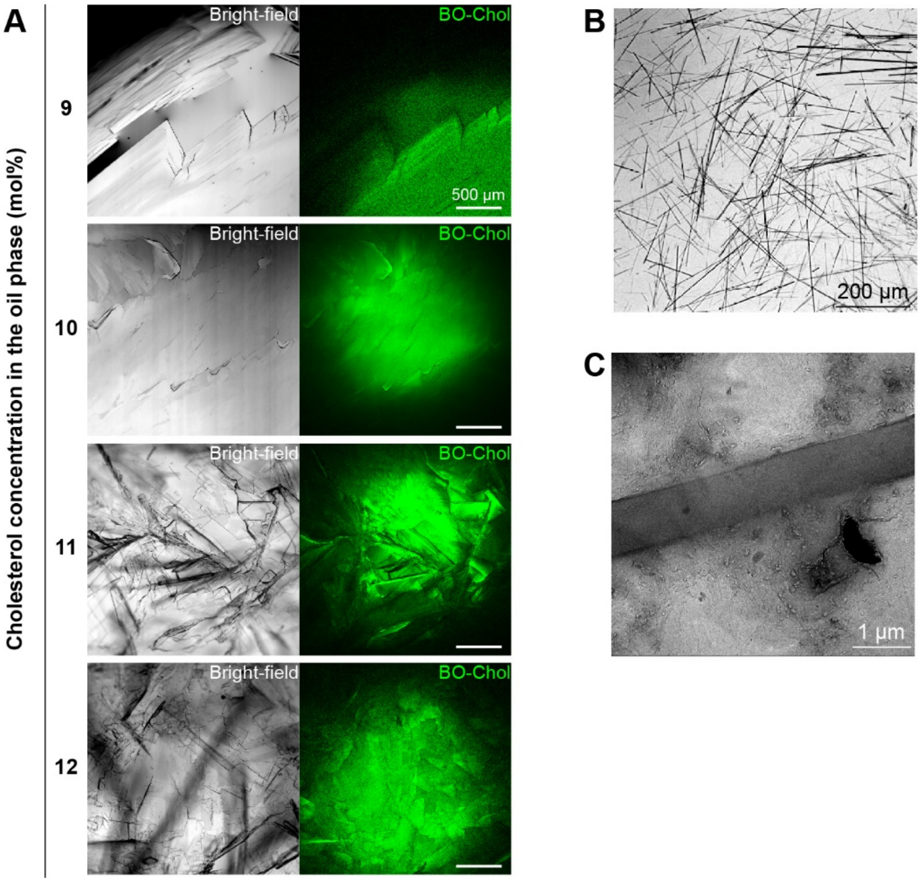


**Fig. S1. Cholesterol crystallization at the oil/water interface and in the oil phase. (A)** Representative images of cholesterol crystals formed at the oil/water interface at the cholesterol concentration in the oil phase of between 9-12 mol%. **(B-C)** Needle-like cholesterol crystals formed in the oil phase were extracted from the oil phase and visualized through the bright-field microscopy **(B)** and cryo-transmission electron microscopy (Cryo-TEM) **(C)**. Cryo-TEM measurements were performed using a Thermo Scientific Glacios Cryo-TEM equipped with a HAADF STEM detector. The microscope was operated at an acceleration voltage of 200 kV, and the samples were maintained at -170°C throughout the measurement. The images were recorded with a defocus value range from -0.5 µm to 1.0 µm.

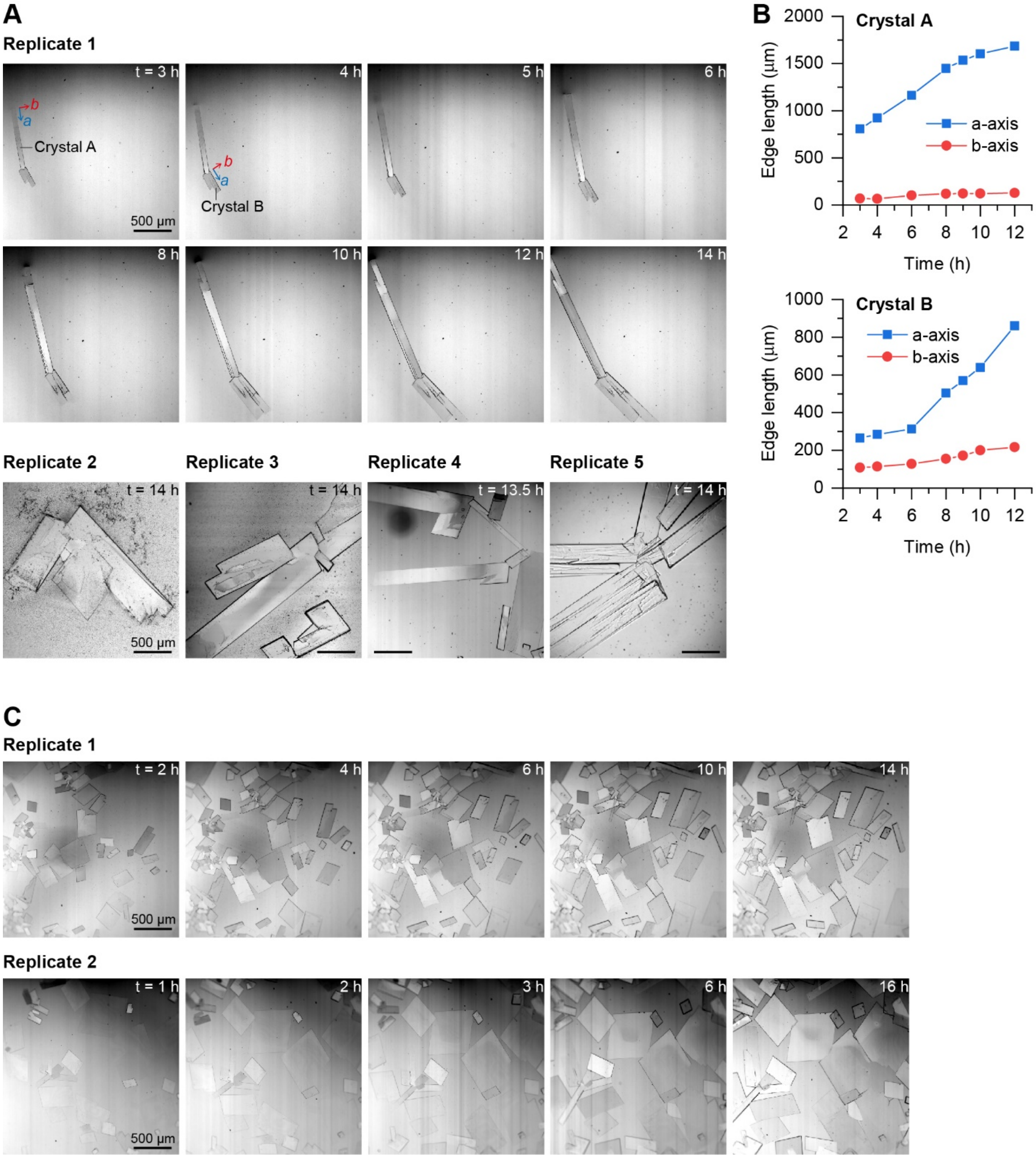


**Fig. S2. Representative images of interfacial cholesterol crystals. (A)** Crystals formed by oil-to-interface adsorption at a cholesterol concentration in the oil phase ($C^{o}_{Chol}$) of 8 mol%. The replicates 2 and 3 are the same crystals shown in Fig. 1E and 2A, respectively. **(B)** The edge lengths of cholesterol crystals (Replicate 1) along the a- and b-axes were measured over time. The crystal growth rate along the a-axis is significantly greater than that along the b-axis, leading to the elongated crystal morphology. **(C)** Crystals formed by MβCD-mediated cholesterol transport at $C^{o}_{Chol}$ of 8 mol%. The replicate 1 is the same crystals shown in Fig. 2B. t represents the time measured from when the oil/water interface was formed.

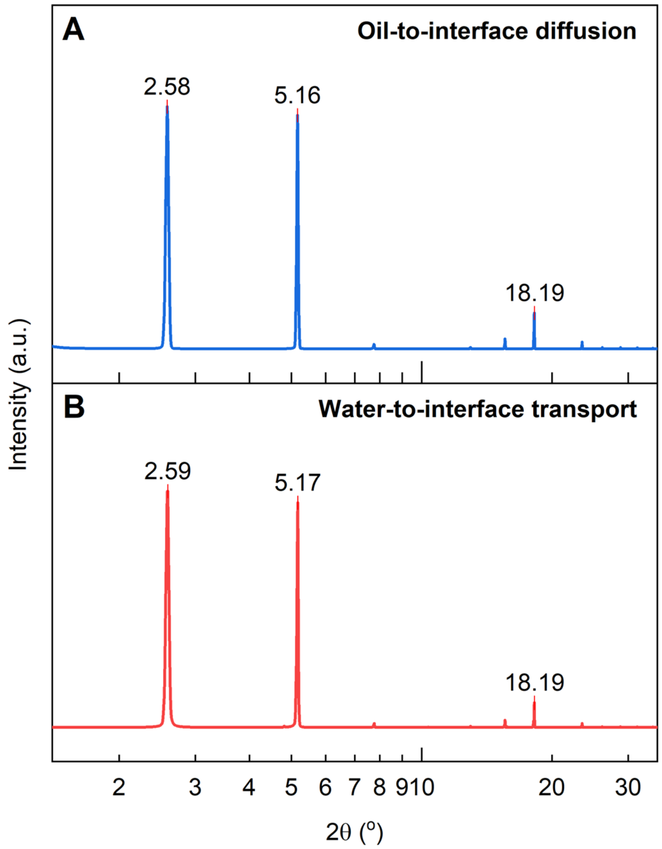


**Fig. S3. X-ray diffraction patterns of cholesterol crystals.** Crystals formed by oil-to-interface diffusion (A) and MβCD-mediated water-to-interface transport (B).

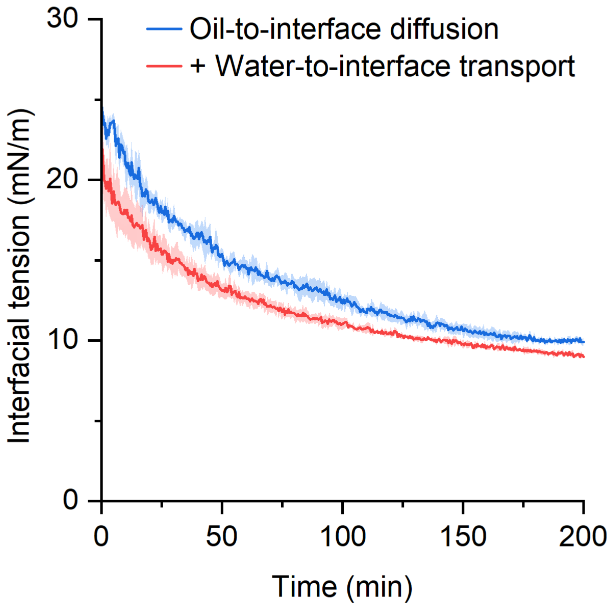


**Fig. S4. Variation in interfacial tension by adsorption of cholesterol at the oil/water interface.** Interfacial tension was quantified using a pendant drop tensiometer, varying according to the different interfacial processes described in the main text. $C^{o}_{Chol}$ was set at 4.5 mol%. The shaded regions represent the standard deviation.

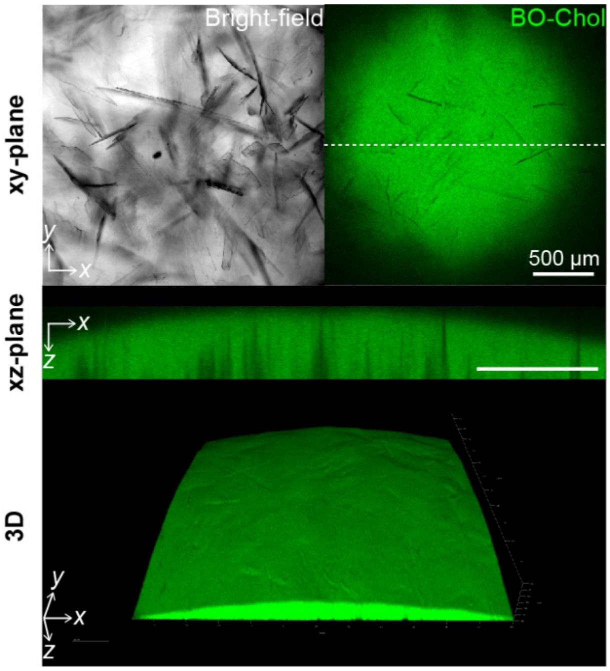


**Fig. S5. Crystallization of cholesteryl palmitate (CP) at the CP concentration in the oil phase of 6 mol%.** CP crystals were visualized through bright-field and BO-Chol fluorescence imaging at the oil/water interface. The xz-plane image corresponds to the cross section at the dotted line of the xy-plane image.

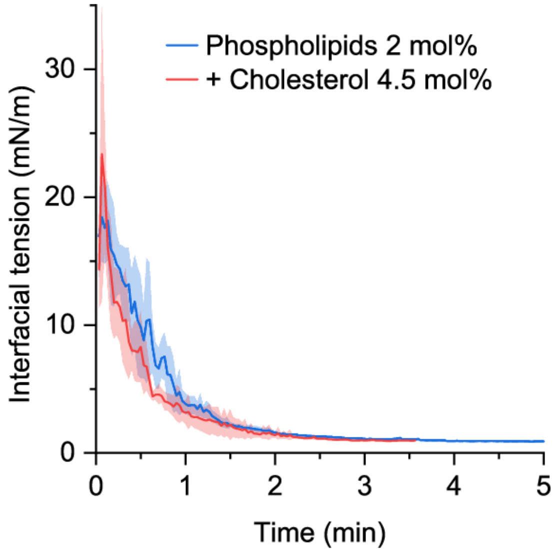


**Fig. S6. Variation in interfacial tension by adsorption of phospholipids and cholesterol at the oil/water interface.** Interfacial tension at the phospholipid-laden interface was assessed using a pendant drop tensiometer. The shaded regions represent the standard deviation.

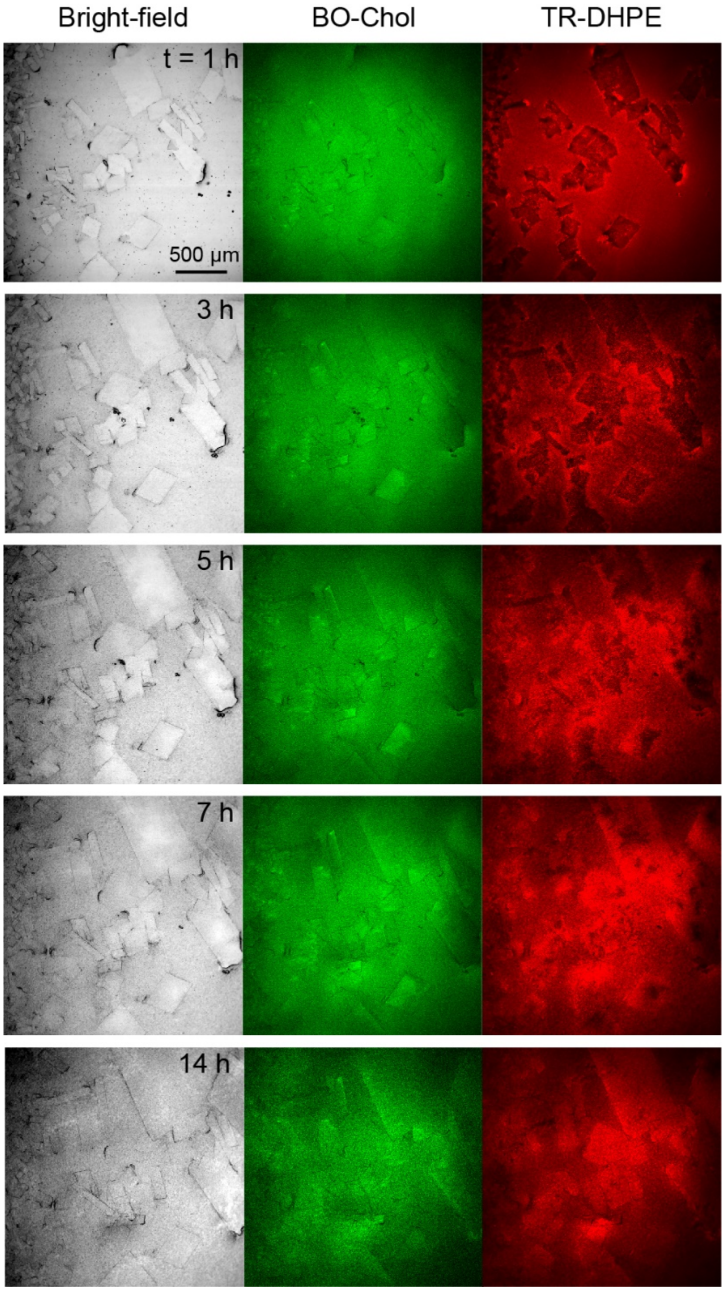


**Fig. S7. Temporal growth of cholesterol crystals at the phospholipid-laden oil/water interface.** The crystals are identical to those shown in Fig. 4C. t represents the time measured from when the oil/water interface was formed.

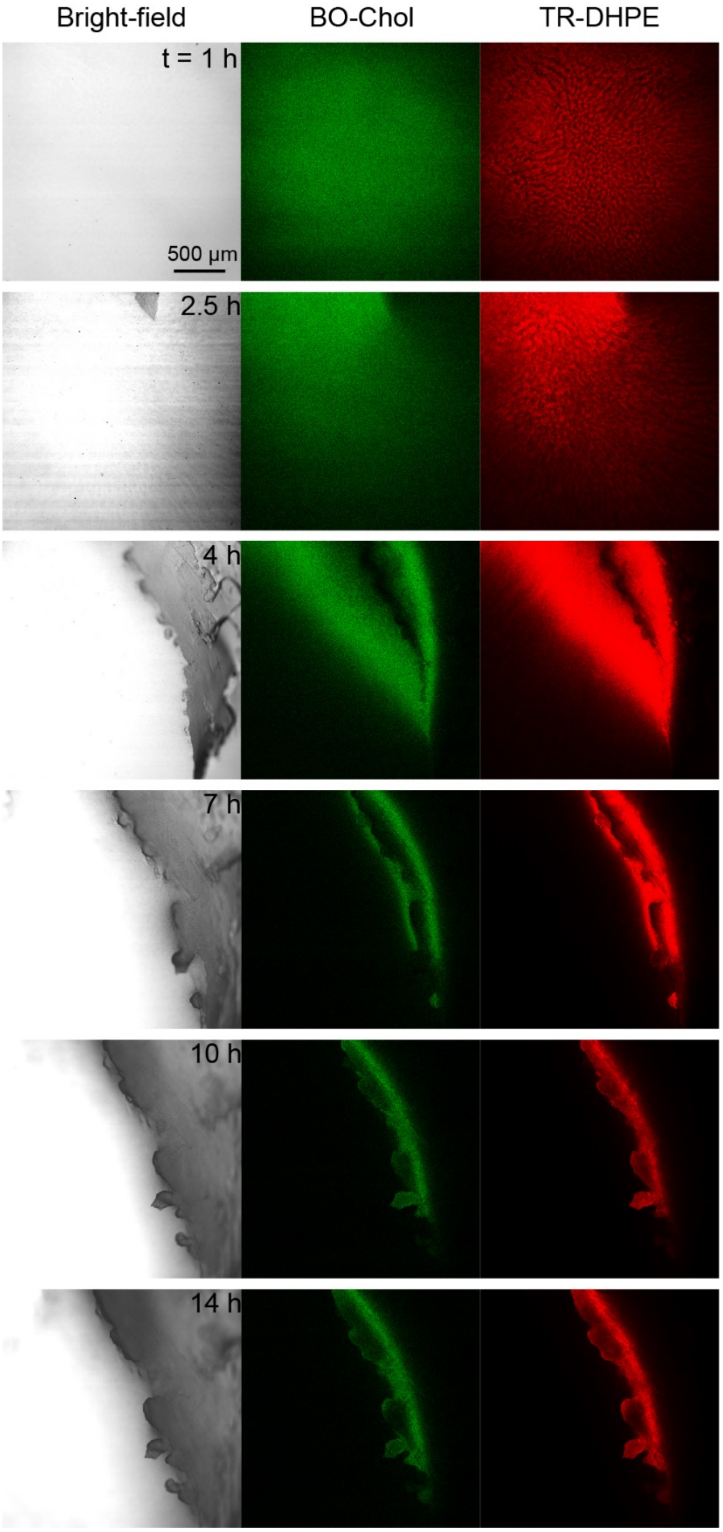


**Fig. S8. Temporal growth of flat, sheet-like cholesterol crystals at the phospholipid-laden oil/water interface.** The crystals are identical to those shown in Fig. 5C. t represents the time measured from when the oil/water interface was formed.

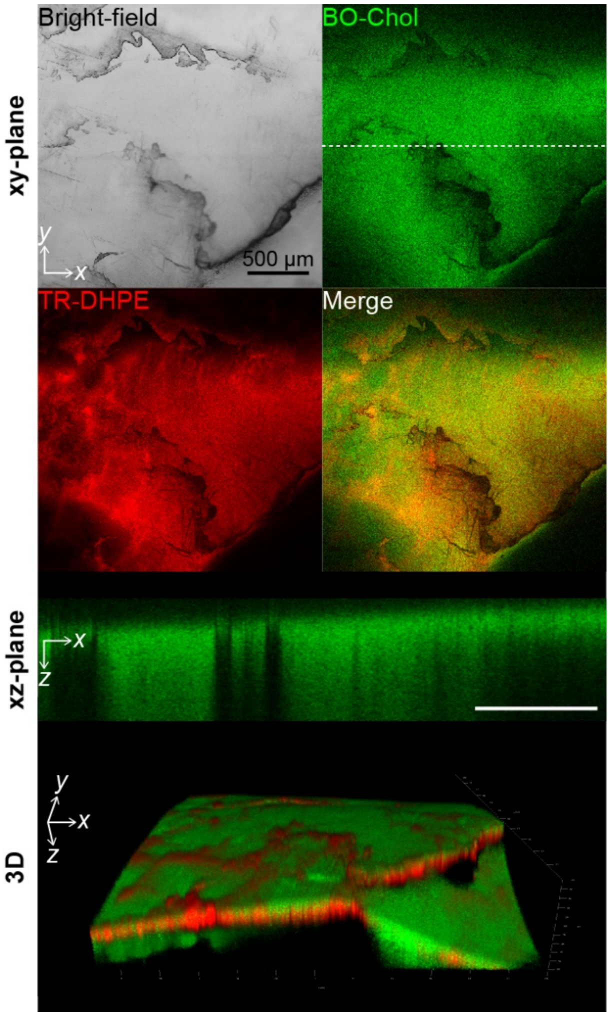


**Fig. S9. Flat, sheet-like cholesterol crystals grown at the phospholipid-laden oil/water interface.** The interfacial cholesterol crystals were visualized by the bright-field and fluorescence imaging at cholesterol/phospholipid concentrations of the oil phase of 13/2 mol%. The xz-plane view represents the cross-section at the dotted line of the xy-plane image.

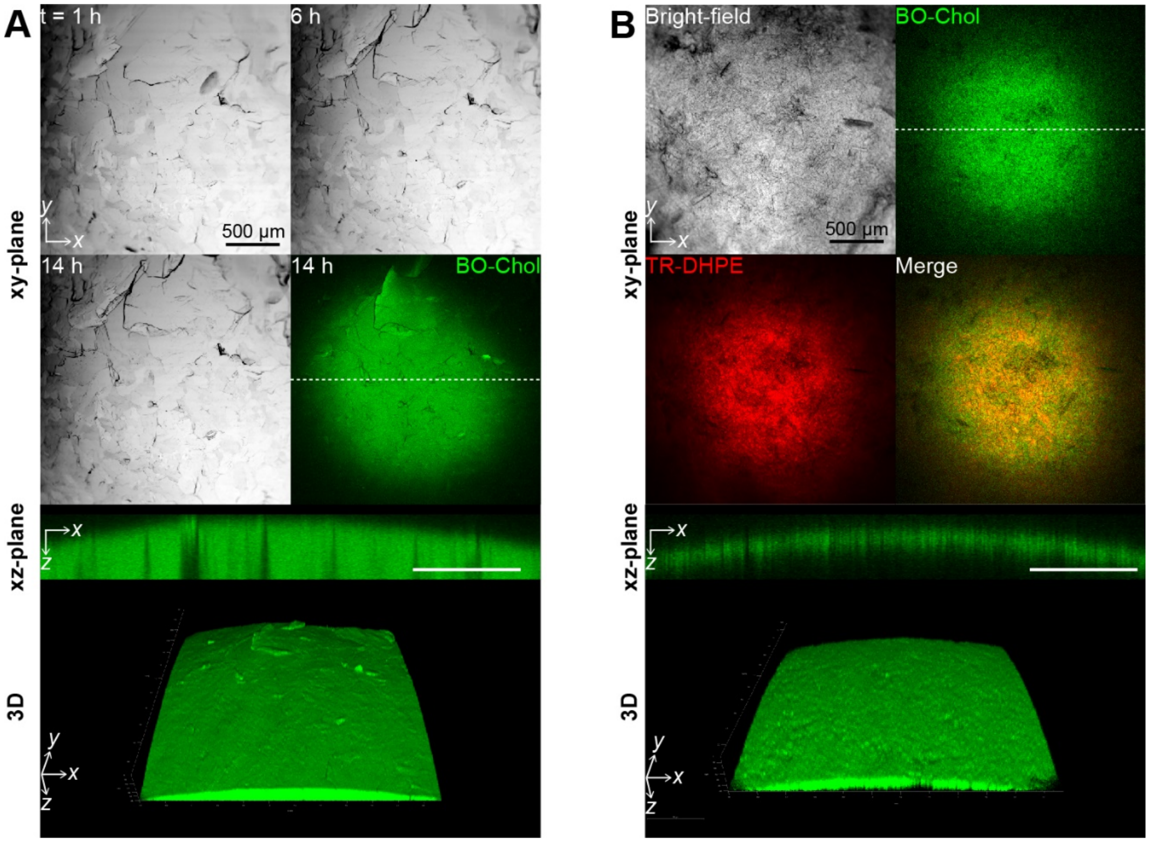


**Fig. S10. The formation of curved, sheet-like cholesterol crystal aggregates at the oil/water interface.** Crystals were visualized through bright-field and BO-Chol fluorescence imaging at cholesterol/phospholipid concentrations of 10/0 mol% **(A)** and 12/2 mol% **(B)** in the presence of MβCD. The xz-plane image represents to the cross section at the dotted line of the xy-plane image. t represents the time measured from when the oil/water interface was formed.

**Table S1. Bragg peaks and D-spacings of cholesterol crystals formed with cholesteryl esters**

| Crystals | 2θ* (°) | D-spacing* (Å) | # of trials (N) |
|---|---|---|---|
| **Cholesterol** | 2.59 ± 0.02 | 34.20 ± 0.11 | 3 |
| | 5.17 ± 0.02 | 17.09 ± 0.02 | 3 |
| | 18.19 ± 0.02 | 4.87 ± 0.00 | 3 |
| **CP** | 1.68 ± 0.00 | 52.65 ± 0.15 | 5 |
| | 3.33 ± 0.01 | 26.52 ± 0.06 | 5 |
| | 4.99 ± 0.01 | 17.70 ± 0.03 | 5 |
| **Cholesterol/CP** | 1.67 ± 0.00 | 52.76 ± 0.14 | 4 |
| | 2.60 ± 0.01 | 34.01 ± 0.11 | 4 |
| | 3.33 ± 0.00 | 26.52 ± 0.03 | 4 |
| | 4.99 ± 0.00 | 17.71 ± 0.02 | 4 |
| | 5.18 ± 0.01 | 17.05 ± 0.04 | 4 |
| | 18.20 ± 0.01 | 4.87 ± 0.00 | 4 |
| **Cholesterol/CO** | 5.17 ± 0.01 | 17.07 ± 0.03 | 3 |

*Expressed as mean ± standard deviation
